# Redox-active molecules in bacterial cultivation media produce photocurrent

**DOI:** 10.1101/2022.12.03.518941

**Authors:** Yaniv Shlosberg, Matthew C. Smith, Nathan. S. Nasseri, Emile J. Morin, Jakkarin Limwongyut, Alex S. Moreland, Guillermo C. Bazan, Andrea S. Carlini

## Abstract

In recent years, global climate change has urged the importance of developing new clean energy technologies that may replace the combustion of fossil fuels and help to reduce carbon emissions. A promising, yet intrinsically complex solution, is the utilization of bacteria as electron donors in microbial fuel cells (MFCs). In this work, we show that just the cultivation media for bacterial growth, which is based on yeast extract, is sufficient for generating electrical current in a bio-electrochemical cell (BEC). We apply cyclic voltammetry and 2-dimensional fluorescence spectroscopy to identify redox active molecules such as NADH, NAD^+^, and flavines that may play key roles in electron donation. Finally, we show that upon illumination, current production is enhanced 2-fold. This photocurrent is generated by a variety of metabolites capable of photochemical reduction, enabling them to donate electrons at the anode of the BEC. Future investigations into the role of culture media versus that of living cells in MFCs may aid in reducing overall production and maintenance costs, and enhance power output performance.

## Introduction

Fossil fuel combustion has been the primary source of energy production around the world. Their staggeringly high carbon emissions and non-renewability risks long-term climate changes and global disasters and has prompted the development of novel clean energy technologies. Among the leading technologies that dominate the renewable energy market are: solar cells^1^, turbines^2^, and hydro plants^3^. A biologically inspired concept for microbial fuel cells (MFCs) was invented over a hundred years ago by Potter et al^4^. This method allows bacterial cells to be integrated with an electrochemical cell and applied as electron donors at the anode to produce clean energy without carbon emission.

Bacteria can perform external electron transport (EET) through two mechanisms: 1) direct electron transfer (DET) ^5–7^ and 2) mediated electron transfer (MET)^8–13^. DET is conducted by transmembranal conductive protein complexes such as metal respiratory complexes ^14,15^ or conductive *pili* ^16–19^. The bacterial species *Shewanella oneidensis* ^17^ and *Geobacter sulfurreducens* ^20^ are rich in these complexes and are able to perform high DET-based exo-electrogenic activity. Natural MET occurs through the secretion of electron donors from cells such as NADH^21^, phenazines, and quinone derivatives ^8–13^. Electrical current production can be enhanced with the addition of exogenous electron mediators^22–27^, the utilization of advanced electrode materials^28,29^, and growing biofilm architecture in close association with the anode^30^.

However, several major challenges persist in translating these technologies for manufacturing such as regulating cell growth, cell waste removal, and environmental regulations^31^. In the case of bacterial-based MFCs, microbial overgrowth can cause clogging or massive cell death which may become an environmental risk; this precludes mass production, system longevity, and ability to produce high electrical power^32^. Furthermore, a continuous supply of cultivation media is required to maintain cell viability in MFCs. This can be addressed by using photosynthetic microorganisms^33–35^ or live macro-organisms (e.g. seaweeds, plant leaves/roots, mammalian fuel cells, and sea-anemones^36–42^) that act as electron donors in bio-electrochemical cells (BECs). Live organisms can act as a continuous source of reducing molecule generation for long term electricity production.

Regardless of the approach, both MFC and BEC development is limited by an inadequate understanding of their underlying energy metabolism pathways^43^. Studies on bio-electricity generation from isolated components such as enzymes^28,44–47^, photosystems^48^, chloroplasts^49^, and thylakoid membranes^50^ introduces a simplified setup for studying photosynthetic organism, but does not provide a scalable or affordable solution. In this work, we show that simply bacterial cultivation media Luria-Bertani (LB) alone, which is comprised of inexpensive yeast extract, consists of reducing molecules sufficient for electrical and photoelectrical current generation. We identify some of these molecules as NADH, NAD^+^, and flavins in our simplified setup.

## Materials and Methods

All chemicals were purchased from Merck, MilliporeSigma, and Fisher Scientific

### LB media preparation

10 g Tryptone, 5 g yeast extract, and 10 g NaCl were dissolved in 1L double distilled water under sterile conditions. Unless otherwise stated, solutions were autoclaved at 120 °C for 30 min.

### Absorbance measurements

Absorption measurements were collected at 750 nm (Nanodrop 2000 UV-Vis spectrophotometer, Thermo Fisher Scientific).

### 2D – Fluorescence microscopy (FM) measurements

The 2D-FM was conducted using a Cary Eclipse fluorimeter (Varian) with excitation and emission slits bands of 5 nm, applying of a voltage of 700 V on the photomultiplier detector.

### Electrochemical Measurements

All electrochemical measurements were conducted on a PalmSens4 potentiostat equipped with a commercially available screen-printed electrodes (SPEs) with graphite counter electrode and silver-coated silver chloride reference electrode, and graphite working electrode. CV experiments, utilized SPEs a 4 mm diameter working electrode (Metrohm, 6.1208.110) within a window of 0 – 1.2 V and a scan rate of 0.1 V/s over 5 cycles per sample. Unless otherwise stated, chronoamperometry experiments, utilized SPEs with a 1 mm diameter working electrode (Basi, SP-1401).

### Chronoamperometry

CA was performed by illumination of each solution with an LED source. For most experiments, a white LED (100 W, 400-700 nm) simulated a broad spectrum solar source. Light intensity at the electrode surface was determined to be 1000 W/m^2^. For selective illumination at 470 nm, a ThorLabs T-cube LED driver and ThorLabs Collimated LED light source were used. The power source was configured to deliver 3.40 W to the LED. The irradiance intensity was measured to be 44.6 W/m^2^ using an OceanOptics USB4000 spectrometer. MetroOhm C110 SPE (Carbon working and counter electrode, Ag reference electrode) were used for all experiments. A standardized drop size of 100 μL was placed on the SPE, ensuring the entire surface of the working, counter, and reference electrodes were covered by the sample. Each measurement was conducted using a brand new electrode to prevent fouling. In all measurements, a potential of 0.9 V was applied on the working electrode. In the case of adding NADH in the middle of the CA measurement. A drop of 20 µL of 6 µM NADH was carefully pipetted on top of the NaCl solution for a final NADH concentration of 2 µM. This gentle placement of a drop on top of another drop was conducted to prevent signal disruption that may occur due to fluctuation in the drop during the spike.

### Time Constant Calculation

Time constants and their fit were calculated by plotting normalized photocurrent rise as a function of time for each light-on cycle, and fitted with an exponential form to yield a time constant of T = 27.5 s to reach 63.7% of a quasi-steady-state value (∂[α(t)]/∂[t]∼0).

### UPLC-MS measurements

All ultra-high performance liquid chromatography mass spectrometry (UPLC-MS) data was collected via a Waters Xevo G2-XS Time-of-Flight mass spectrometer with electrospray ionization, monitored at 254 nm. Additional detection wavelengths corresponding to the maximum absorption of some metabolites (340 nm for NADH and 450 nm for FAD) were also assessed in tandem. The flow rate was kept constant at 0.25 mL/min. The column temperature was 40 °C. Samples injections were 5 µL each. Gradient separation of quinone derivatives was achieved with Method 1, and detection of NADH, NAD^+^, and flavines utilized Method 2 (see **Table 2**).

Benzoquinone (BQ) and 2,6-dimethoxybenzoquinone (DMBQ) were derivatized with 2,4-dinitrophenylhydrazine (DNPH) to amplify signal detection of target quinones, based on an assay reported by Stagge et al^62^. Briefly, 36 mg of DNPH was dissolved in a 15 mL solution of 12 M HCl, H_2_O, and ACN (2:5:1). LB media (100 µL) was mixed with this DNPH solution (1500 µL), and incubated 18 hr at room temperature before being diluted to 5 mL with ACN. As positive control for DMBQ involved spiking LB media with 15 µL DMBQ (0.89 mM) standard. For NADH, NAD^+^, and flavine detection, LB samples and controls were buffered in a 10 mM ammonium acetate buffer adjusted to a pH of 9.

### NADH/NAD^+^ Quantification Assay

LB samples that were autoclaved or not autoclaved were subjected to quantification of NADH/NAD^+^ using an NAD^+^/NADH Colorimetric Assay Kit (ab65348, Abcam, Cambridge, United Kingdom). Autoclaved samples were measured before and after either chronoamperometry or cycle voltammetry treatment. Absorbance kinetics were collected on a SpectraMax iD5 plate reader.

## Results and Discussion

### Identification of redox-active molecules in Luria-Bertani media

Previous studies have shown redox-active molecules such as NADH^21^ and flavines^17^ may play a major role in mediating electron transfer between live organisms and the anode of BECs to produce current. These molecules can be selectively identified in complex matrixes based on their unique 2D-fluorescence spectral fingerprints^51^. To determine whether such molecules exist in LB, we applied 2D-fluorescence maps (λ_Ex_ = 230 – 600 nm, λ_Em_ = 250 – 800 nm) (**Fig. 1a**). These spectra showed characteristic fluorescence fingerprints for NAD^+^ (λ_Ex_ = 300 nm, λ_Em_ = 450 nm), NADH (λ_Ex_ = 360 nm, λ_Em_ = 450 nm), and flavins (λ_Ex_ = 450 nm, λ_Em_ = 510 nm)^21,51^. The NADH concentration in LB (1.78 µM) was determined empirically using a calibration curve of fluorescence intensity (λ_Ex_ = 350 nm, λ_Em_ = 450 nm) at increasing NADH concentrations ranging from 0 to 4 µM (**Fig. S1**).

**Fig.1.**
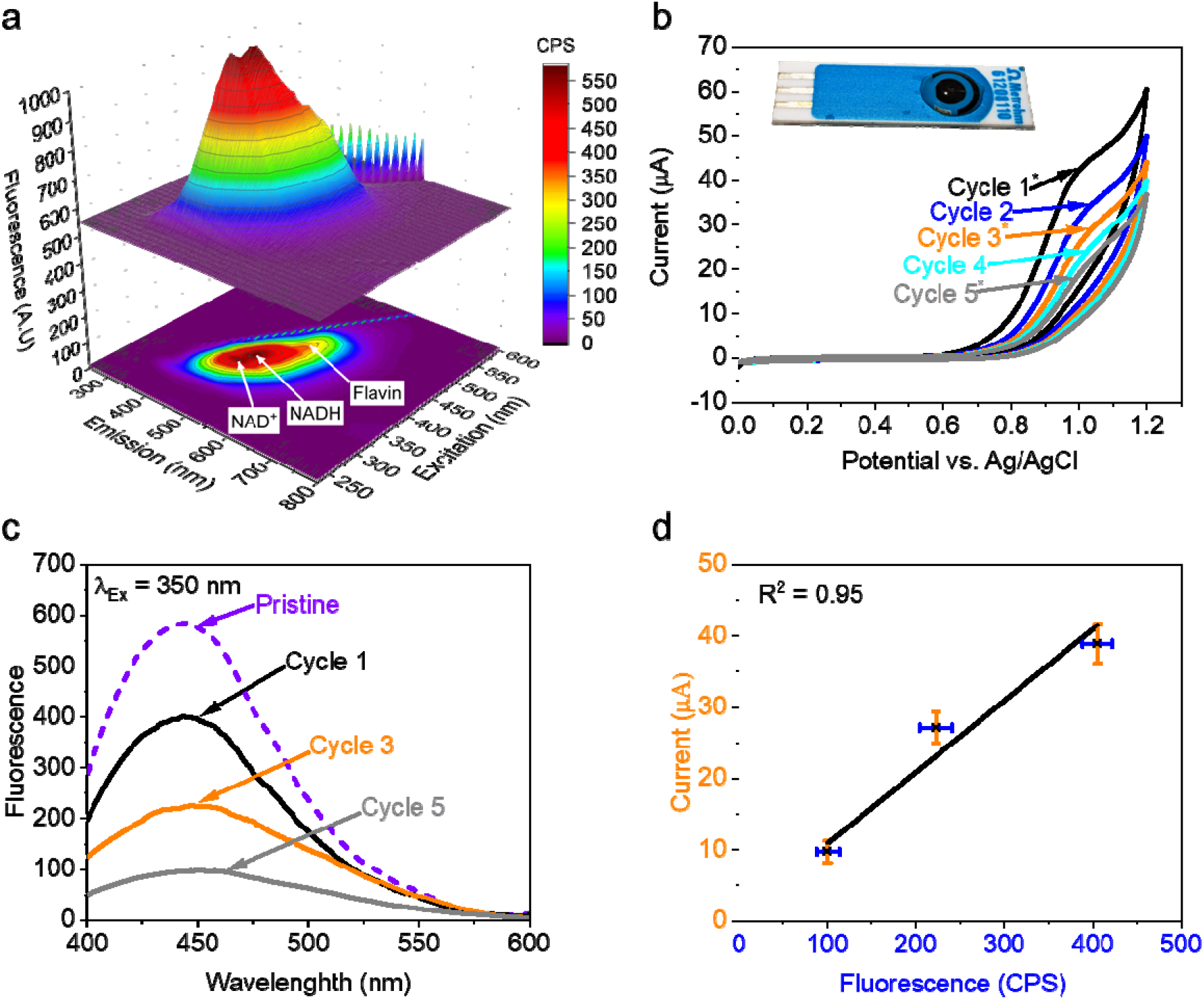
Identification of redox-active molecules and electric potentials in Luria-Bertani media. **a** 2D - fluorescence spectra of LB (λ_Ex_ = 230 – 600 nm, λ_Em_ = 250 – 800 nm) shows the presence of NAD^+^, NADH, and flavines. **b** Redox activity of LB constituents analysed by CV (0.0 – 1.2 V, scan rate 0.1 V/s, n = 5 cycles), showing the fingerprint of NADH. Cycles 1-5 (black, blue, orange, cyan, and grey, respectively). Asterisks mark the cycles numbers in which aliquots were collected for fluorescence measurements Inset shows an optical image of LB media (50 µL) suspended over a commercially available SPE. (**c-d**). **c** Fluorescence spectra (λ_Ex_ = 350 nm, λ_Em_ = 400 -600 nm) of LB pristine (purple) and after 1, 3, and 5 CV cycles (black, orange, and grey) shows a reduction in signal associated with NADH oxidation at 0.9 V. **d,** Maximal fluorescence intensity (λ_Em_= 450 nm) after 1, 3, and 5 CV cycles as a function of current intensities of the same cycles at 0.9 V. The linear fit shows an R-squared value of 0.95. Values are mean ± SD (n = 3 repetitions).

To further study the identity of the redox-active species in LB, cyclic voltammetry (CV) measurements were conducted using screen printed electrodes (Methrohm 61208110) with a graphite anode and cathode and Ag coated with AgCl reference electrode, 5 cycles were scanned (0 -1.2 V, scan rate = 0.1 V/s) (**Fig. 1b**). Voltammograms show a peak with a maximum at 0.9 V, which we correlate with the fingerprint of NADH^41^ At this potential, NADH can be oxidized at the anode. To assess this, the fluorescence spectra (λ_Ex_ = 300 nm, λ_Em_ = 450 nm) of LB after 1, 3 and 5 cycles of CV was measured (**Fig. 1c**), whereby the decrease in current intensities from CV directly correlated with a decrease in fluorescence emission at 440-470 nm (λ_Ex_ = 350 nm) (**Fig. 1d**). A recovery assay showed that addition of exogenous NADH (2 µM) after multiple cycles of CV recapitulated the signal at 0.9 V (**Fig. S2**). Based on the identification of NADH in LB, we postulated that it could be a key molecule for producing electrical current in a BEC.

### Luria-Bertani media produces current and photocurrent in a BECs

Based on the observation that LB consists of redox-active molecules, we wished to estimate its ability to act as an electron donor in a BEC (**Fig. 2**). Furthermore, we sought that among a variety of components in the LB, there may be molecules that can produce photocurrent. To address these questions, chronoamperometry (CA) measurements of LB were performed under dark or illumination conditions (**Fig. 2a** and **Fig. S3**). A potential of 0.9 V was applied to the anode, corresponding to the observed peak potential from CV measurements (**Fig. 1**). While a higher potential could have increased current generation, we wished to avoid unforeseen production of peroxides (1.2 V) and other electrochemical byproducts. Continuous illumination of LB media for 30 min was conducted using a white light emitting diode (LED) at 1000 W/m^2^ and CA was measured (**Fig. 2b**). A maximal photocurrent of ~ 0.7 µA/cm^2^ was obtained, which was stable during the entire measurement (**Fig. 2b** and **Fig. 2c**). Using a similar electrochemical setup, our measured photocurrent density (0.6 µA/cm^2^) is approximately five times greater than that produced by isolated *E. coli* in PBS, which was initially cultivated in LB media (**Table 1**) ^52^. Additionally, this output is comparable in magnitude to photocurrent produced by several species of cyanobacteria, which relies on photosynthesis to generate electrical current. To eliminate the possibility that the current generation was conducted by direct absorbance of light by the anode, we measured photocurrent production with an optically transparent control solution of NaCl 0.1 M. No photocurrent was produced, verifying illumination of the electrodes does not affect our measurements. We attribute the ∼0.5 µA/cm^2^ dark current density from NaCl to the oxygen evolution reaction (OER) that occurs at 0.9 V^53^. Photocurrent generation from LB was measured in dark/light intervals of 100 s each, showing an enhancement of ∼ 0.7 µA/cm^2^ in generated current density during illumination conditions (**Fig. 2d** and **Fig. 2e**). We observe a gradual drop-off in baseline current density associated with dark conditions. We suggest this decrease originates from the consumption of NADH and other possible reducing molecules over time, as was shown in the CV measurements. Despite an initial reduction in photocurrent magnitude during each illumination cycle, the rate of photocurrent generation is unaltered, with a time constant of 27.5 s (**Fig. 2f**).

**Fig.2.**
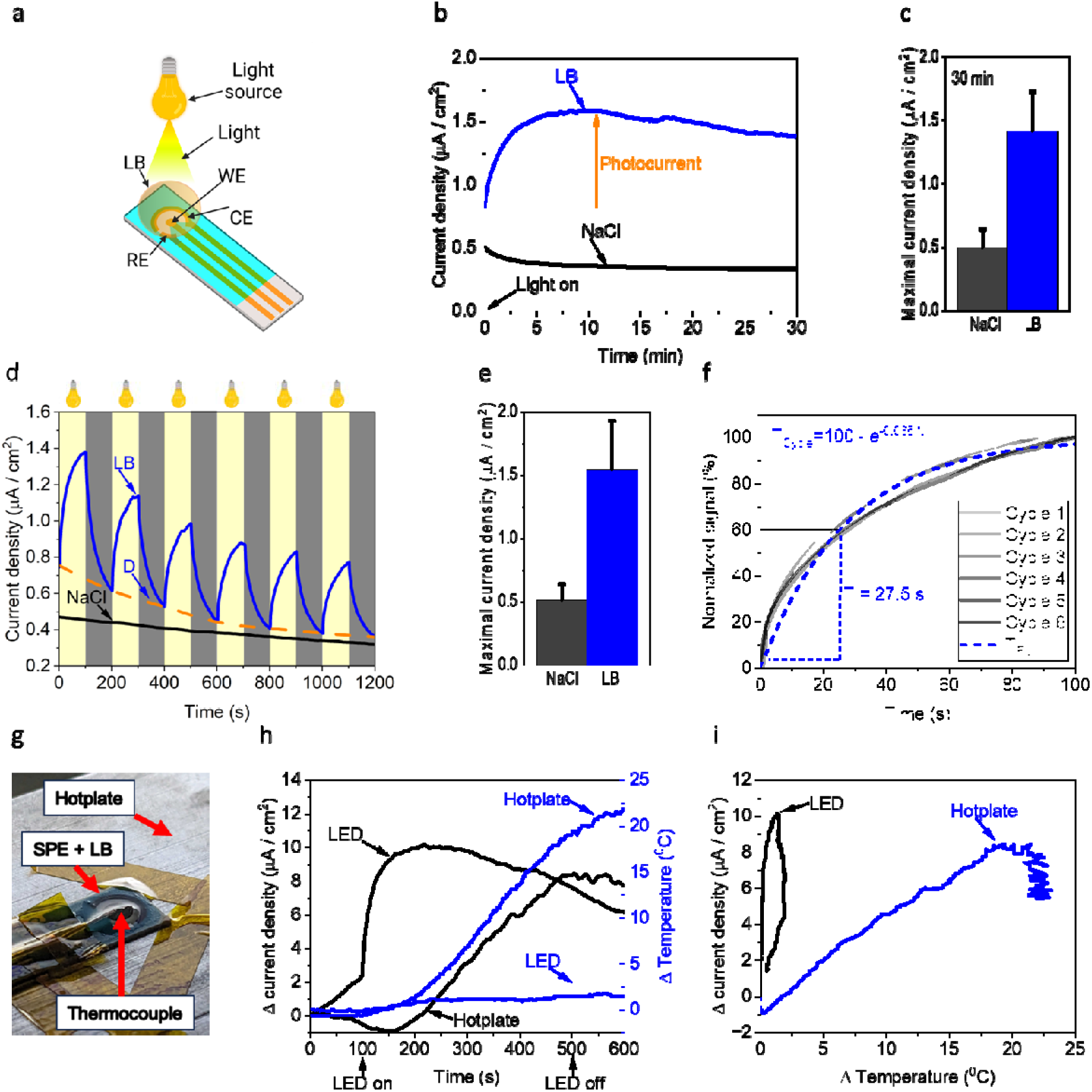
Luria-Bertani media produces dark and photo current in a bio-fuel cell. CA and temperature of LB and NaCl 0.1 M were measured in dark and under white light irradiation or in dark upon hotplate heating. A potential of 0.9 V was applied at the working electrode. **a**. Schematic of the system. A drop of 50 µL LB was placed on screen printed electrodes. Light irradiation (Intensity at the electrodes surface = 1000 W/m^2^) was conducted from above using a light bulb. Black arrows show the working, counter and reference electrodes (WE, CE, and RE respectively). **b**. CA measurements of NaCl (black), and LB (blue) under continuous illumination for 30 min. **c**. Maximal current density of NaCl and LB over 30 min of continued illumination. (n = 3 ± SD**) d**. CA measurements of NaCl (black), and LB (blue) over 6 light/dark illumination cycles. An orange dashed line represents the dark current based on an extrapolation of the current values before the turn on of the light in each interval. Grey and yellow panels with a light bulb icon on top represent dark and light conditions, respectively. **e**. Maximal current density of NaCl and LB for light/dark illumination cycles (n = 3 samples ± SD, 6 cycles each). **f**. Plot of normalized photocurrent rise as a function of time for each light-on cycle, and fitted exponential with an average time constant of T = 27.5 s to reach 63.7% of a quasi-steady-state value. **g** Schematic of the system while using a hotplate and a thermocouple sensor. **h** Current density (black) and temperature changes (blue) of LB upon white LED irradiation or a heating treatment using a hotplate. **i** A correlation between current density and temperature changes under white LED irradiation (black) or hotplate heating (blue).

**Table 1.** A comparison of photocurrent generation from LB and microorganisms in BECs.

| Electron donor | Class | Working electrode | Potential vs Ag/AgCl | Max current | Ref |
| --- | --- | --- | --- | --- | --- |
| *Escherichia coli* | Bacteria | Graphite | 0.9 V | 0.6 $\mu\text{A} / \text{cm}^2$ | 52 |
| *Synechocystis*<br>sp.6083 | Cyanobacteria | Graphite | 0.5 V | 0.5 $\mu\text{A} / \text{cm}^2$ | 21 |
| *Acaryochloris marina*<br>MBIC 11017 | Cyanobacteria | Graphite | 0.5 V | 0.3 $\mu\text{A} / \text{cm}^2$ | 21 |
| *Synechococcus elongatus*<br>PCC 7942 | Cyanobacteria | Graphite | 0.5 V | 0.3 $\mu\text{A} / \text{cm}^2$ | 21 |
| *Dunaliella*<br>*Salina* | Microalgae | Graphite | 0.5 V | 2.5 $\mu\text{A} / \text{cm}^2$ | 38 |
| *Trichodesmium*<br>*Erythraeum* | Cyanobacteria | Graphite | 0.5 V | 3.5 $\mu\text{A} / \text{cm}^2$ | 59 |
| LB | Bacterial cultivation media | Graphite | 0.9 V | 0.7 $\mu\text{A} / \text{cm}^2$ | This work |

**Table 2.**
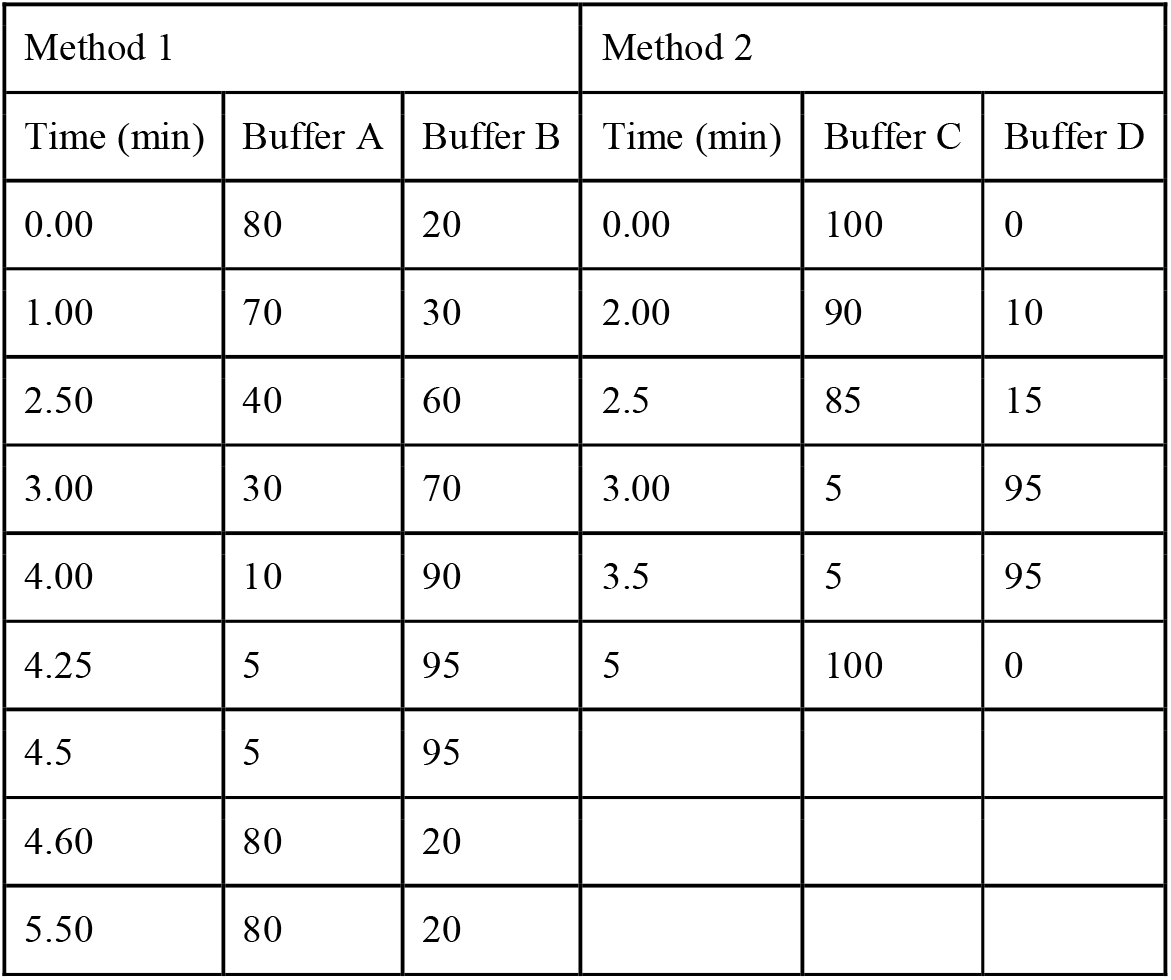
UPLC/MS solvent systems in this study. Method 1, for the detection of quinone species, uses H_2_O with 0.1% formic acid (Buffer A) and ACN with 0.1% formic acid (Buffer B). Method 2, for the detection of flavins (NAD/NADH and FAD/FADH_2_), uses 10 mM ammonium acetate with 0.1% formic acid (Buffer C) and ACN with 0.1% formic acid (Buffer D).

To validate that the photocurrent increase is not a consequence of heat generated by illumination, a four-channel thermoprobe measured the temperatures of ambient air in the room, surface of a dry anode, inside an NaCl or LB droplet. Continuous illumination showed similar rises in solution temperatures of 4-4.5 ^0^C (**Fig. S4**). As the NaCl control showed no photocurrent production in **Fig. 2b**, we conclude that photocurrent produced in LB media is not derived from these temperature changes. To further support this conclusion and decouple the influence of heat on current production, LB was analyzed on SPEs (Metrohm, 6.1208.110 with an WE area = 0.11 cm^2^) on a hotplate in the dark and compared with samples irradiated with a white LED for 600 s (**Fig. 2g-i**). Despite excessive heating from the hotplate (Δ+23.1 ^0^C), the maximal current production (Δ+8.4 µA/cm^2^) was lower than that produced with the LED light (Δ+10.2 µA/cm^2^). Notably, LED irradiation only elevated the temperature by 2.0 ^0^C (**Fig. 2h**). A correlation between the photocurrent generation and the temperature change (**Fig. 2i**) shows that photocurrent production from the LED is decoupled from its minor heating effects, strengthening our conclusion that direct light absorption at the anode is not the major cause for the photocurrent production by LB. Notably, while using the SPE with the WE area of 0.11 cm^2^, the produced current density under LED irradiation was about 6 times greater than previously obtained while using SPEs with a WE of 1 mm (**Fig. 2b**). We postulated that this difference my originate from a different surface tension of the SPE, resulting in a lower contact angle for the LB droplet. This increased wettability and decreased droplet height is expected to accelerate electrode reactivity and enhance light penetration at the WE-solution interface, respectively^54^.

### NADH plays a major role in current production in dark but not in light

As the spectral fingerprint of NADH and NAD^+^ was obtained in the 2D-fluorescence spectra of LB (**Fig. 1a**), we wished to analyse their individual contributions to photocurrent production. To account for potential overestimation of each fluorophore caused by overlap in their fluorescence signatures in a mixed solution, we attempted analytical methods for quantification. Ultra performance liquid chromatography mass spectrometry (UPLC-MS) yielded no quantifiable detection of NADH or NAD^+^. This agrees with literature reports of ionization suppression in mixed component solutions, on-source fragmentation during mass spectral characterization, and the lability of fragile bonds in these metabolites that result in a variety of decomposition products^55^. We thus relied on a quantitative colorimetric enzyme-based assay kit to measure the ratios of each species before and after oxidative treatment by CV and CA (**Fig. 3a**). Initially, samples show nominal levels of NAD^+^, with the dominant species being NADH (**Fig. 3b**). We reveal a significant rise in NAD^+^ following 5 cycles of CV, whereas an even larger conversion is observed following CA treatment at an applied potential of 0.9 V under constant illumination for 30 min. We note that in the absence of autoclaving to sterilize our LB media, the concentrations of NADH are ∼ 3 times higher. We next examined the detrimental effect of autoclaving on photocurrent density and observe no significant difference (**Fig. 3c**). This supports our rationale that photocurrent generation under illumination is not dominated by NADH. To strengthen this conclusion, we studied the dark current and photocurrent production by pure NADH and NAD^+^ (**Fig. 3d** and **Fig. 3e**). CA of NaCl was conducted in the dark, followed by addition of NADH (2 µM final concentration) at 100 s. This caused a rapid spike, which we attribute to a momentarily high concentration gradient experienced prior to solvent mixing. Rapid equilibration after ∼15 seconds yields an elevated current density of ∼ 1.4 µA / cm^2^ attributable to dark current from NADH. This dark current is likely a large component of what we observe in **Fig. 2d**. Illumination of the sample at 200 s showed a nominal current increase of about 0.01 µA/cm^2^, followed by a steady decline. We suspect that this initial current increase may result from the photocatalytic activity of nicotinamide adenine dinucleotide that may directly donate electrons at the anode, or reduce oxygen molecules generated by oxygen evolution reaction at 0.9 V to O_2_.^-^ that mediates electrons to the anode ^56^. The gradual decrease in current can be explained by continuous consumption of NADH at the anode. The resulting NAD^+^ species, which cannot be oxidated at the anode, was next studied for photocurrent production. The photocurrent generated with 2 µM of NAD^+^ in NaCl was ∼0.06 µA / cm^2^, which is an overestimate of the levels present in autoclaved LB. Nonetheless, this study demonstrates that increasing consumption of NADH results in the generation of a photoactive molecule.

**Fig. 3.**
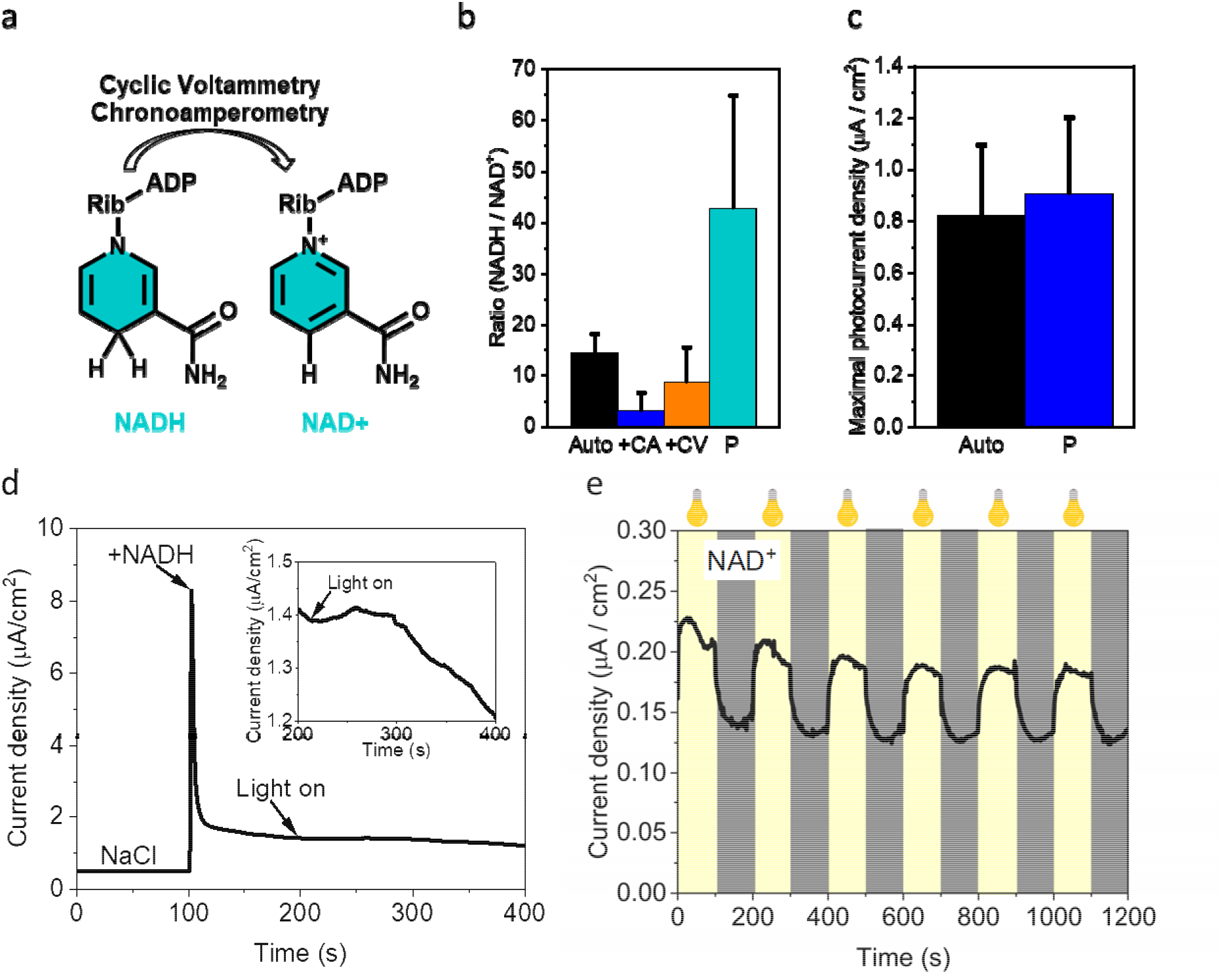
NADH plays a major role in current production in dark but not in light. The concentration ratio of NADH/NAD^+^ was determined using a colorimetric kit of NADH before and after CA and CV electrical treatment or autoclaving. CA of pristine and autoclaved NADH. Photocurrent generation was measured for NADH and NAD^+^. **a** A schematic description of the oxidation of NADH to NAD^+^ by CA and CV treatments. **b** NADH/NAD^+^ concentration ratio after autoclaving (Auto, black), After 30 min of CA (blue), and CV (orange), and of pristine (without any treatment) (P, cyan). (n = 3 ± SD) **c** Maximal photocurrent generation by non-autoclaved (black) and autoclaved (blue) LB. (n = 3 ± SD) **d.** CA of NaCl with addition of NADH after 100 s in dark, light was turned on after 200 s. **e** CA of NAD^+^ measured in dark/light intervals of 100 s. The grey and yellow stripes with the bule icon represent dark and light respectively.

### Benzoquinones are not involved in the photocurrent production

While LB is composed of yeast extract, it consists of numerous molecules that originate from the internal content of yeast such as various proteins, lipids, metabolites, and more. For example, previous studies about MFCs, showed that the electron transfer mediation between the bacterial cytoplasm and the anode outside of the cell can be conducted by various quinones^57^. To detect benzoquinones in LB media, benzoquinone (BQ) and 2,6-dimethoxy benzoquinone (DMBQ) were derivatized with 2,4-dinitrophenylhydrazine (DNPH). UPLC-MS measurements reveal the presence of BQ ([M+H]^+^ = 289 m/z), but not of DMBQ (**Fig. S5**). Yet, as the only major peak obtained in the CV was NADH, we suggest that the contribution of benzoquinones in LB to the current production is minor. This is supported by CA of BQ, which produced a small photocurrent of 0.016 µA / cm^2^ (**Fig. S6**).

### Flavins in LB are major player in the photocurrent production

Previous studies have reported photo-electrochemical reactions converting FAD into FADH_2_ and other forms of FAD or FADH radicals (**Fig. 4a**), which are oxidized at the anode of a BEC to produce electrical current^58^. As the fingerprint of flavines was detected in the 2D-fluorescence measurements (**Fig. 1**), we wished to assess their potential contribution to photocurrent generation in LB media. Similarly with NADH and NAD^+^, UPLC-MS detected no significant amounts of these major metabolites, however, a specific degradation fragment of FAD, lumichrome, was identified in trace amounts (**Fig. S7**). We posit that ion suppression and partial thermal degradation of the full metabolites in autoclaved LB limit quantification of individual flavins observed by fluorescence. As such, we chose FAD as a model metabolite to study. CA of FAD (2 µM) was conducted under light/dark cycles of 100 s. (**Fig. 4b**). Results showed a maximal photocurrent production of ∼ 0.1 µA/cm^2^ corresponding to ∼ 14 % of the total photocurrent generated by LB. To strengthen the assessment that flavins play an independent role in photocurrent production, we sought to selectively irradiate LB media with a narrow spectrum blue LED at 470 nm, which overlaps with the absorption maximum for FAD, but not NADH and other optically active low wavelength components in LB (**Fig. 4c**). The measurements were conducted on screen printed electrodes which were irradiated from above with an LED_470_ (**Fig. 4d** and **Fig. 4e**). CA of LB under continuous irradiation for 30 min, yielded maximal photocurrents of ∼ 0.1 µA / cm^2^, respectively (**Fig. 4f** and **Fig. 4g**). An irradiation of 2 µM FAD has generated a photocurrent of ∼ 0.015 -/+ 0.004 µA / cm^2^ (**Fig. S8**). Based on this, we conclude that FAD can play a minor role in photocurrent production by LB.

**Fig. 4.**
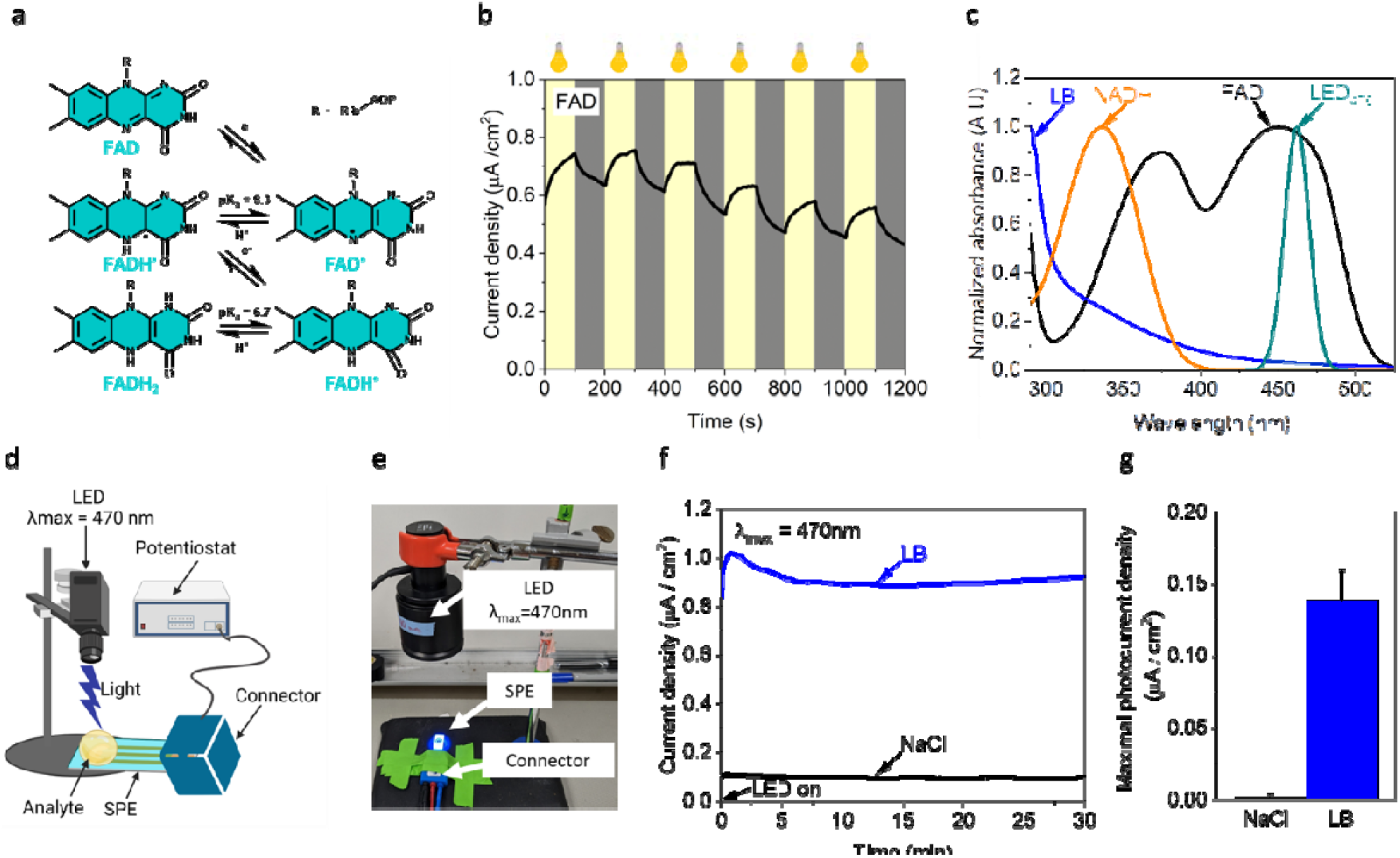
Flavines play a role in the photocurrent production of LB. Absorption spectra of LB NADH and FAD were measured. Photocurrent of LB and FAD were measured under irradiation of white light, and LED_470_ which emits close to the maxima of FAD. **a** A schematic description of the photoelectrochemical reactions of FAD under irradiation. **b** CA of FAD under white light irradiation. **c** Normalized absorption spectra of FAD (black), LB (blue), NADH (orange), and the irradiation spectra of LED_470_ (cyan). **d, e** A schematic description and a photo of the BEC system using screen printed electrodes irradiated by LEDs from top. **f** CA of NaCl (black), and LB (blue) irradiated by LED_470_ for 30 min. **g** Maximal photocurrent generated by NaCl, and LB irradiated by LED_470_. The error bars show the standard deviation over 3 independent repetitions.

### Suggesting a model for the current and photocurrent production

We suggest that in addition to NADH and flavins that were detected in the LB solution, it is likely that many unknown components such as redox-active proteins, vitamins^59^, and scavenging antioxidants (e.g. glutathione)^60^ likely contribute to the dark current or photocurrent produced, herein (**Fig. 5**).

**Fig.5.**
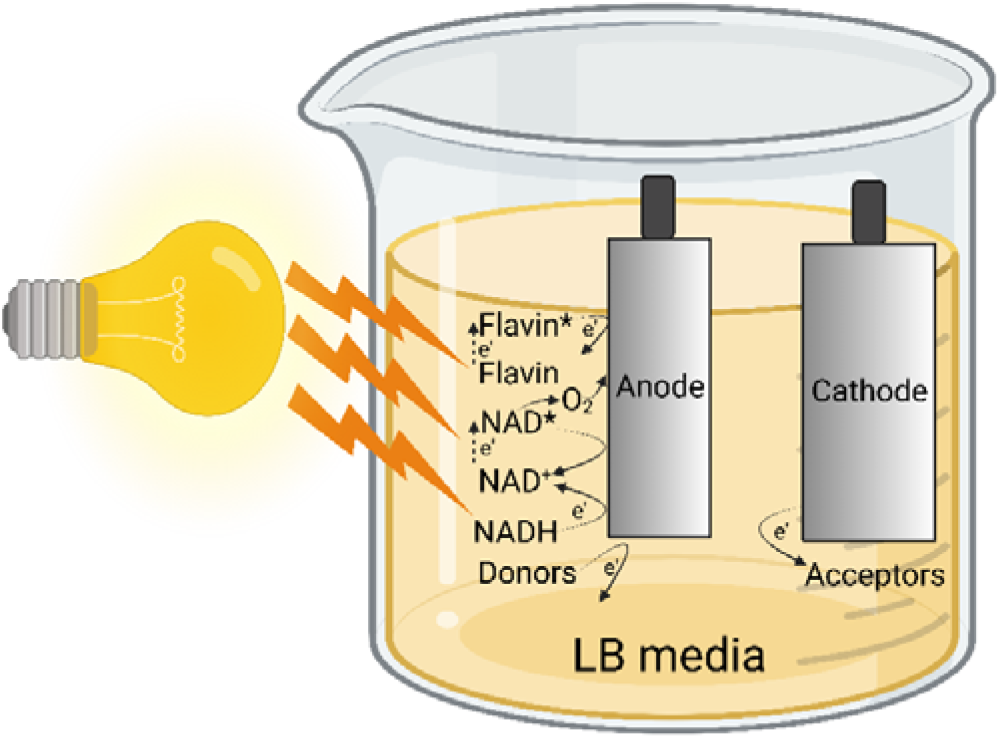
Suggesting a model for the dark current and photocurrent production. NADH, flavins, and other possible electron donors that originate from yeast extract donate electrons at the anode of an electrochemical cell. These electrons flow to the cathode and finally to an electron acceptor that may also originate from the yeast extract component in the LB. Upon light irradiation, flavins, NAD^+^, and NADH may donate electrons via its excited state to the anode. In the case of NAD^+^, the electron transfer may also be mediated by O_2_ molecules formed by the oxygen evolution reaction that occurs at the applied potential of 0.9 V. Flavins may be reduced by photochemical reactions which can be oxidized at the anode.

### Conclusions

In this work, we show that the bacterial cultivation media LB consists of reducing molecules that can apply to produce current and photocurrent in a BEC. We applied fluorescence, colorimetric assays and electrochemical measurements to identify some of the reducing molecules to be NADH and flavins and benzoquinones. The idea that bacterial electron donors in MFCs can be simply replaced by their cultivation media may be a basis for affordable and scalable light-activated BECs that are not hampered by cell death, overgrowth, and nutrient/waste regulation.

## Acknowledgments

We thank the University of California and a UCSB Faculty Research Grant for financial support. Yaniv Shlosberg is supported by the Otis Williams Fellowship. Fluorescence measurements in this study were obtained using central facilities at the UCSB Materials Research Laboratory (MRL), along with technical support from Jaya Nolt. The MRL Shared Experimental Facilities are supported by the MRSEC Program of the NSF under Award No. DMR 1720256; a member of the NSF-funded Materials Research Facilities Network (www.mrfn.org). Some of the figures were prepared using Biorender.com. Thanks to Natasha Cao for manuscript revisions.

## Author contributions

Y.S and A.S.C conceived the idea. Y.S and A.S.C designed the experiments. Y.S performed the main experiments. E.J.M, J.L, A.S.M., M.C.S, and N.S.N assisted in performing the experiments. Y.S and A.S.C wrote the manuscript. YS, G.C.B and A.S.C supervised the research project.

## Supporting Information

**Fig. S1.**
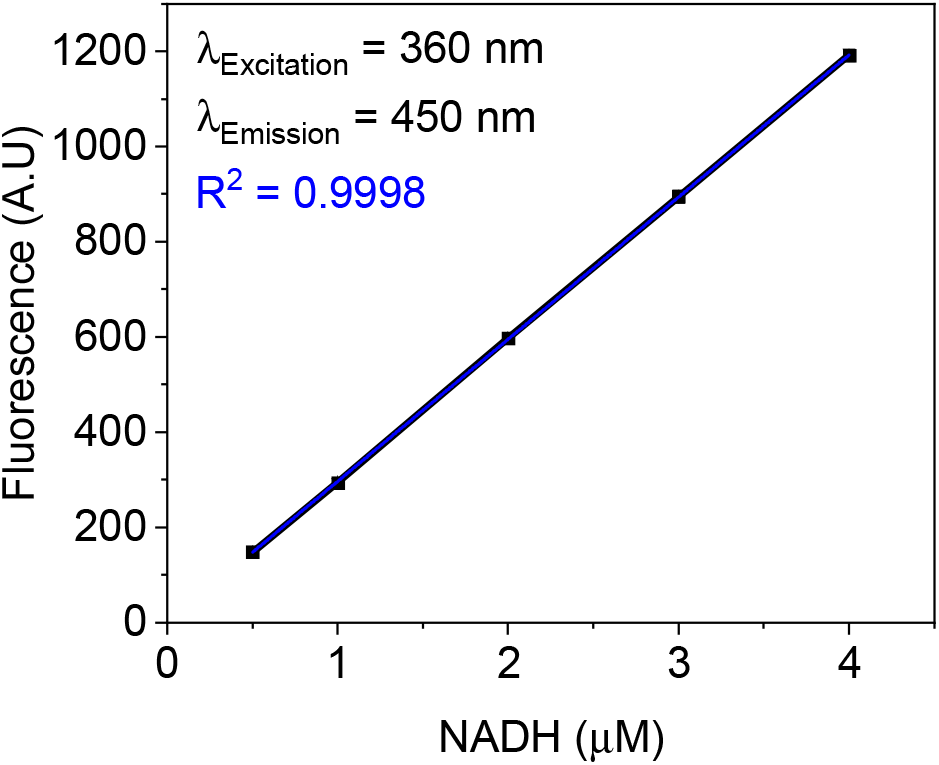
NADH calibration curve. The fluorescence intensity (λ Excitation = 360 nm, λ Emission = 450 nm) of increasing NADH concentrations of (0.5, 1, 2, 3, and 4 µM) was measured. A calibration curve of NADH concentration vs. Fluorescence intensity (black). A linear fit (blue).

**Fig. S2.**
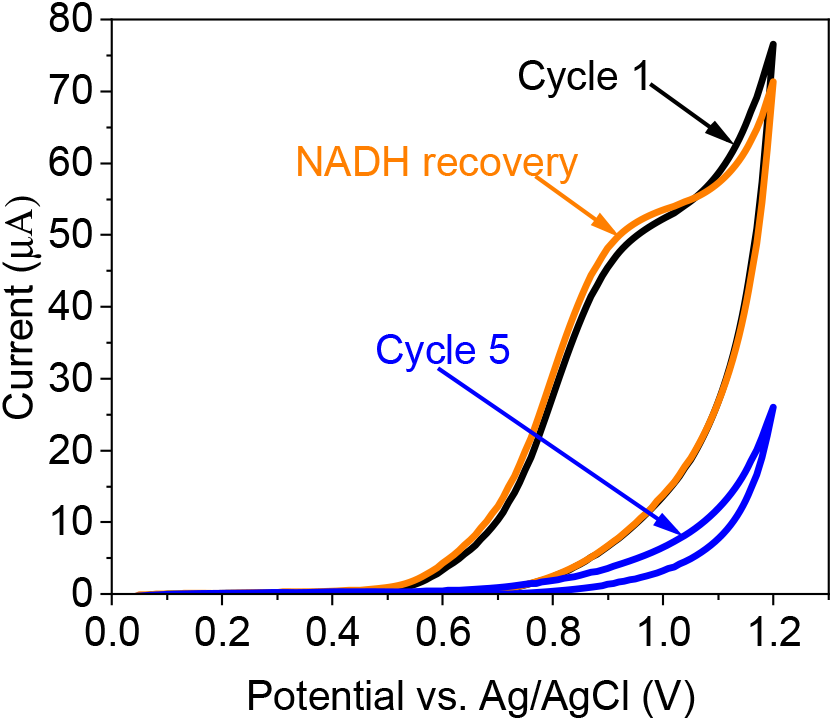
Signal recovery by addition of NADH. CV of LB was measured for 5 cycles to fully oxidize the NADH in the LB. 50 µL of NADH solution was added to the analyte to form a final concentration of 2 µM NADH, and the CV was measured again. Cycle 1 and 5 of LB without the addition of NADH (black and blue). CV of LB after 5 cycles + addition of NADH (orange).

**Fig. S3.**
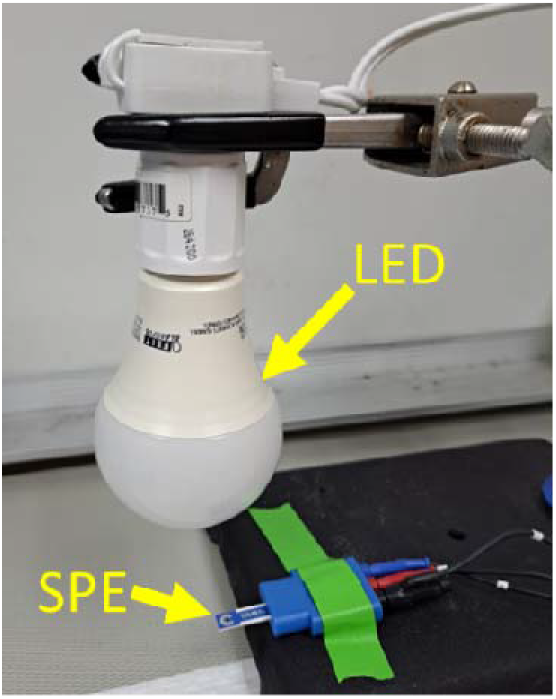
A picture of the measurement system. Yellow arrows mark the LED light source and the SPE.

**Fig. S4.**
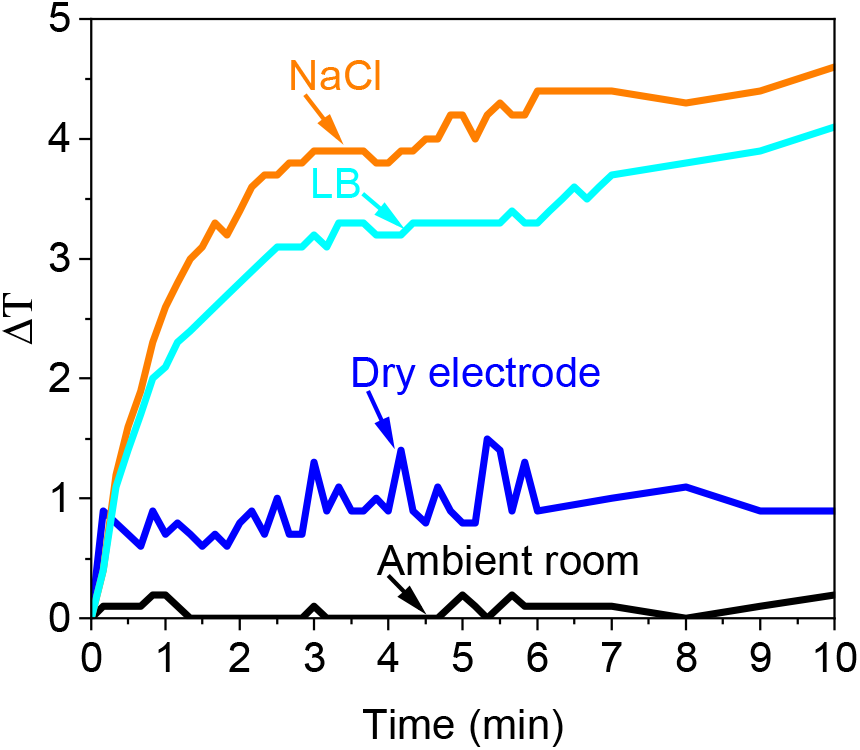
Temperature change of SPE and analytes upon LED irradiation. The temperature change under LED irradiation of the ambient room (black), dry SPE (blue), NaCl (orange) and LB (cyan) drops were measured using a thermocouple sensor.

**Fig. S5.**
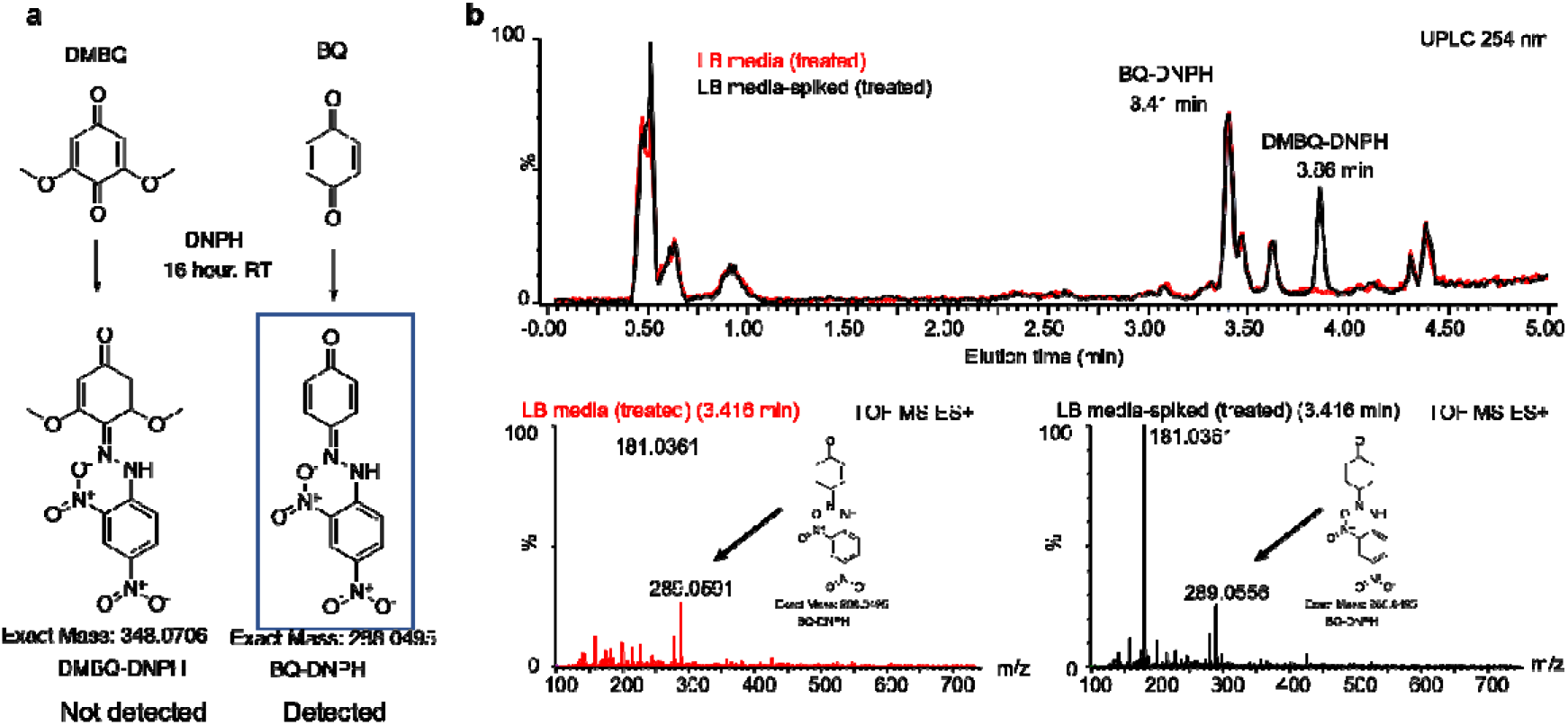
Identification of benzoquinones in LB media by UPLC/MS through signal amplification with DNPH. a) Chemical structures for benzoquinone (BQ) and 2,6-dimethoxybenzoquinone (DMBQ) treated with 2,5-dinitrophenylhydrazine (DNPH) to form charged complexes DMBQ-DNPH (not detected) and BQ-DNPH (detected). **b)** UPLC/MS chromatogram of DNPH-treated LB media with (black) and without (red) DMBQ spike. The eluted peak at 3.41 minutes in both spectra show a mass of 349 m/z, corresponding to the [M+H]^+^ species of BQ-DNPH. 181 m/z corresponds to a degradation product for this complex, as DNPH with a fragment loss of [•OH]. The retention times and detected masses are labelled for each of the peaks.

**Fig. S6.**
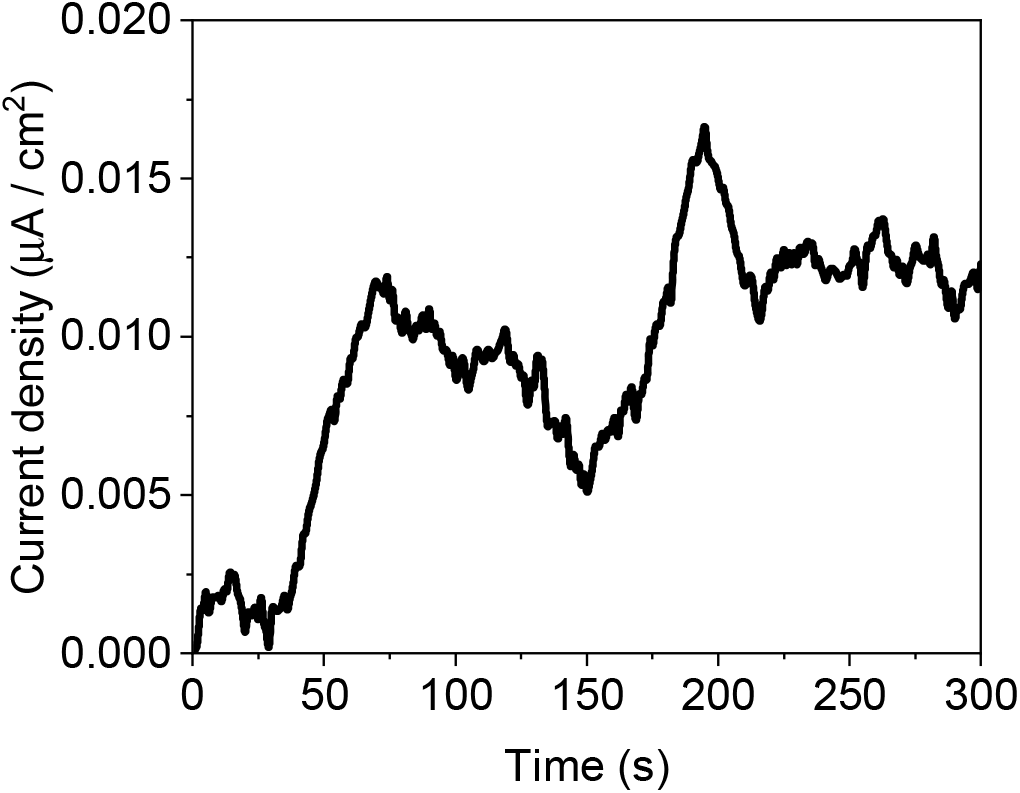
Photocurrent production from BQ. CA of BQ was measured with an applied potential of 0.9 V at the anode under white light irradiation. The current of PBS measured at the same conditions was subtracted from the photocurrent of the BQ. The current at the beginning of the measurements before turning on the light (time = 0 s) was normalized to 0.

**Fig. S7.**
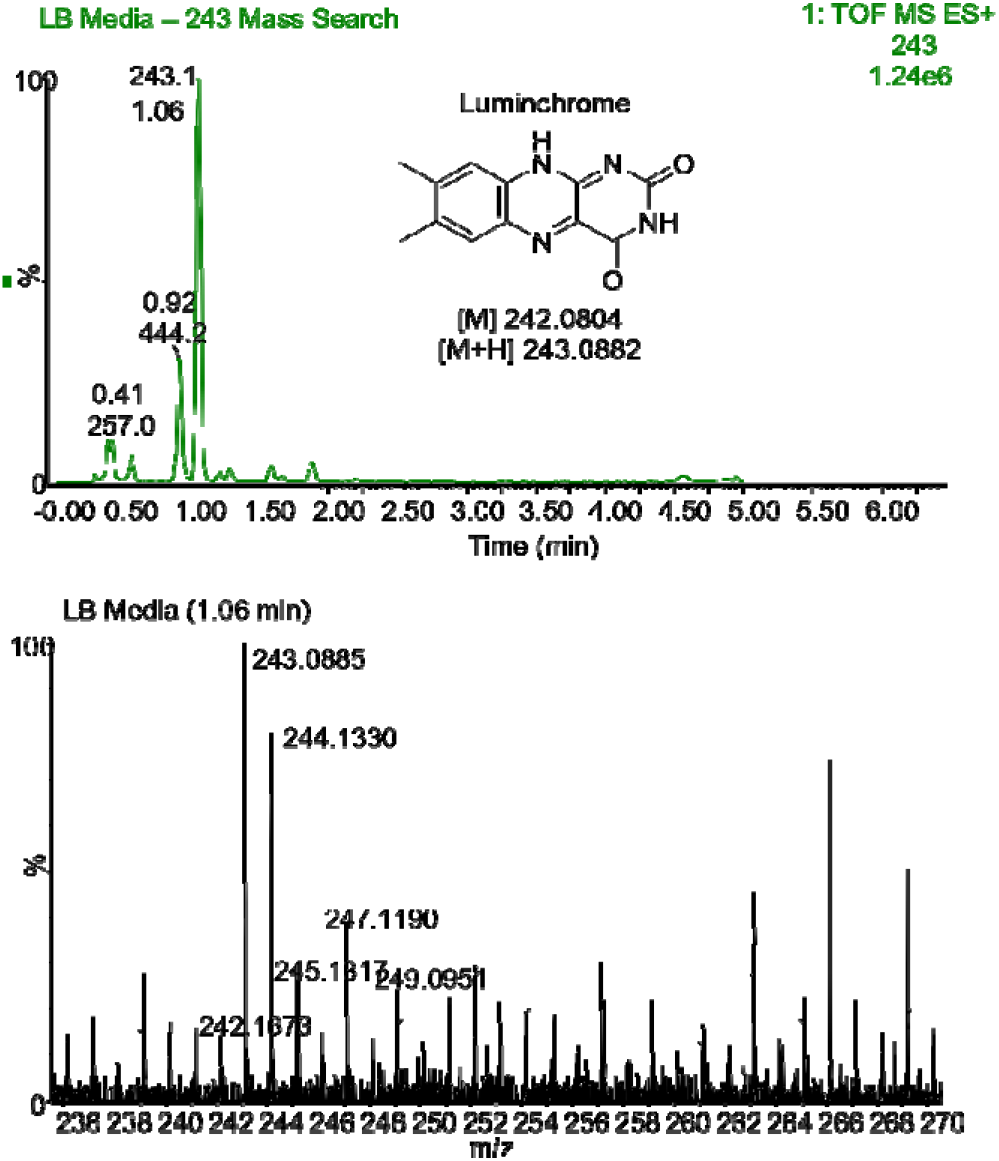
Identification of Lumichrome in LB. UPLC/MS chromatogram of LB media. Specific mass search (top) for 243 m/z shows a major peak for 243.09 m/z at 1.06 min elution time, corresponding to the [M+H]^+^ species for lumichrome with 1.23 ppm error.

**Fig. S8.**
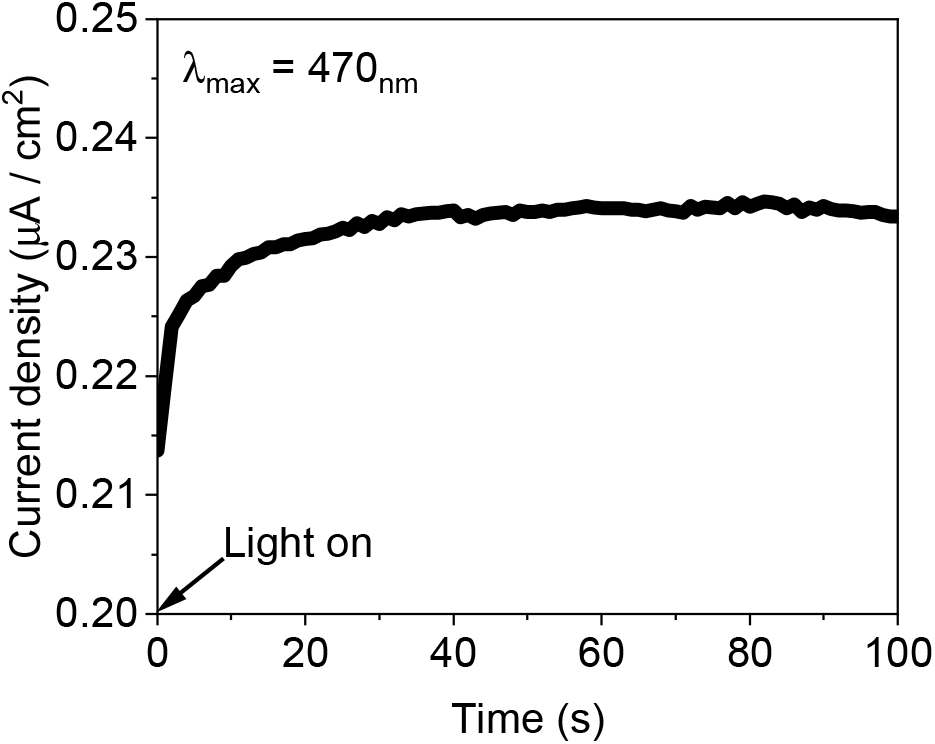
Photocurrent production of FAD under LED_470nm_ illumination. CA of 2 µM FAD was measured under LED_470_nm illumination. **a** a representative CA measurement of FAD.

